# Tailored machine learning models for functional RNA detection in genome-wide screens

**DOI:** 10.1101/2022.09.01.506220

**Authors:** Christopher Klapproth, Siegfried Zöztsche, Felix Kühnl, Jörg Fallmann, Peter F. Stadler, Sven Findeiß

## Abstract

The *in silico* prediction of non-coding and protein-coding genetic loci is an area of research that has gathered large attention in the field of comparative genomics. In the last decade, much effort has been made to investigate numerous properties of nucleotide sequences that hint at their biological role in the cell. We present here a software framework for the alignment-based training, evaluation and application of machine learning models with user-defined parameters. Instead of focusing on the one-size-fits-all approach of pervasive *in silico* annotation pipelines, we offer a framework for the structured generation and evaluation of models based on arbitrary features and input data, focusing on stable and explainable results. Furthermore, we showcase the usage of our software package in a full-genome screen of *Drosophila melanogaster* and evaluate our results against the well-known but much less flexible program `RNAz`.

## 1. Introduction

Experimental and theoretical work in molecular biology is often based on the correct annotation of genomic data and therefore dependent on the ongoing compilation and curation of accessible sequence databases. In the last two decades, large efforts have been undertaken to improve genome 5 annotations. The demand for high-quality genomes led to the emergence of high-throughput sequencing (HTS) techniques and huge amounts of data since the 2000s [1, 2]. Together with the ever increasing available computational power, automated annotation pipelines have supplemented time consuming, error-prone and hardly reproducible manual curation techniques [3]. Still, until recently, annotation efforts have largely focused on protein-coding genes (reviewed by Mudge and io Harrow [4]). However, it has been noted that less than 25% of the transcribed human genes account for the entirety of the 19 000 protein-coding RNAs [5]. Large bodies of research demonstrate that so-called non-coding RNAs (ncRNAs) perform an at least equally diverse range of biological and regulatory functions [6, 7]. Therefore, non-coding RNAs have become a main target of research in molecular biology. Of particular interest to the medical research community is that certain i5 types of ncRNAs have been shown to play an important role in the genesis of cancer and other pathophysiological processes [8].

The ncRNAs known today can be grouped by their functions and mechanisms of action. Some of the most prominent examples are microRNAs, siRNAs, snoRNAs and tRNAs [9]. These molecules display a wide range of features such as folding and assembling into complex superstructures, interactions with other RNAs, DNA or proteins, and the regulation of their activity [10]. Non-coding RNAs have also been found to play a major role in chromatin remodeling and epigenetics [11]. Besides their regulatory function, ncRNAs are often associated with structure-related functions, as for example the 16S rRNA strand working as a scaffold in the 50S subunit of the ribosome [12]. Despite their obviously high prevalence and biological relevance, identification and annotation of ncRNAs is still lagging behind and and incomplete in comparison to protein-coding genes. This is in part due to the popularity of poly-A enrichment protocols for HTS in mutual agreement with a higher interest in protein-coding genes in medical studies, which have been in the center of attention since the advent of computational biology [13]. The result is a far greater number of established methods for protein-coding gene prediction.

A subset of well-described non-coding RNAs are characterized by a significant degree of secondary structure conservation, which is often being associated with biological function, as indicated in Figure 1 and discussed, e. g., by Santosh et al. [15] and Rana [16]. For hammerhead ribozymes, the fact that function is tightly coupled to specific structural elements, while large parts of the sequence are almost freely adaptable is readily being used as constraint for the design of target-specific ribozymes [17].

**Figure 1:**
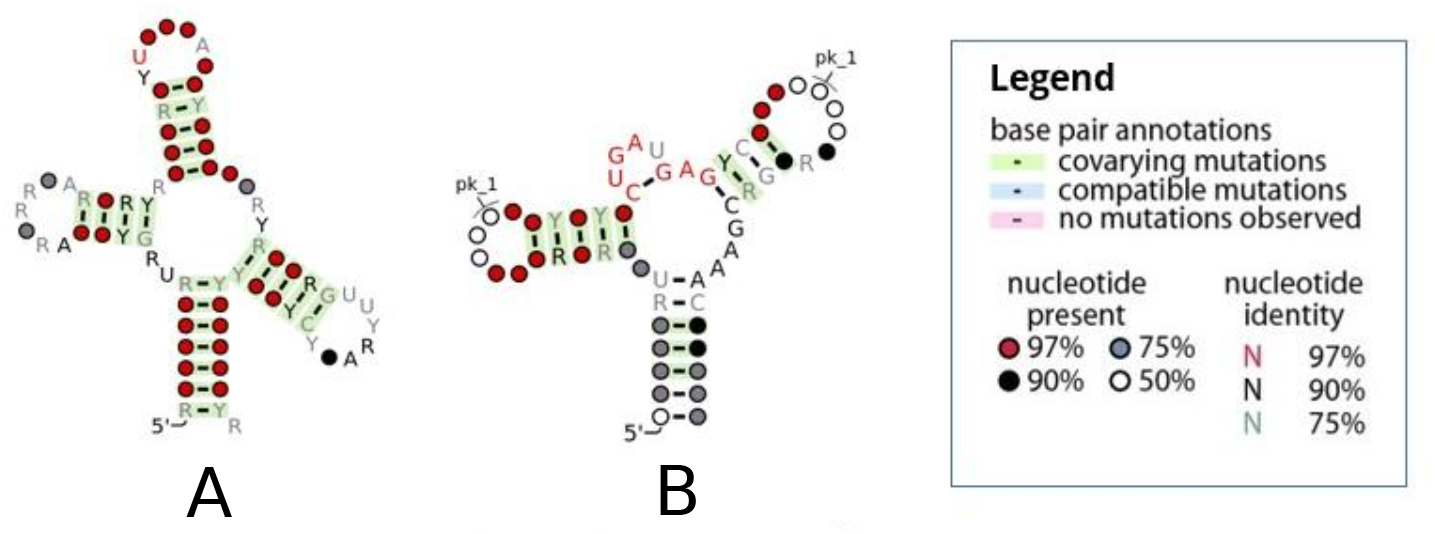
Secondary structure is a common indicator for biological function in the annotation of non-coding RNAs. Many examples, such as the depicted tRNA (A) and Hammerhead Ribozyme (B), show a particularly high degree of conservation of a common structure and a strong evolutionary selection towards mutations that stabilize it. The high degree of observed covarying and compatible mutations strongly support the hypothesis that their structure is required for the sequence to retain its structure-to-function relationship. Figures were generated using `R2R` [14].

It follows that evolutionary conservation of RNA secondary structure is considered a promising predictor of function in the *de novo* identification of ncRNA elements. In this context, a common approach is the prediction of the most stable secondary structure as the one exhibiting the minimum free energy (MFE) fold for a given sequence [18]. Both MFE and intrinsic properties of nucleotide sequences are used to achieve an approximate classification of a given sequence as non-coding or other by tools such as `RNAz, Evofold` and others [19, 20].

The majority of recently developed automatic annotation pipelines focus on the classification of confirmed transcripts, e. g. from high-throughput RNA sequencing experiments, as non-coding RNAs or, more common, long non-coding RNAs (lncRNAs) against a background of proteincoding sequences. Prime examples of such tools include `CPAT, CPC2` and `CNCI` [21, 22, 23]. This approach, however, has the limitation of classifying ncRNAs by classifying all other transcripts as protein-coding, instead of assuming an ambiguous background. The limitations of such a two-way classification approach for the identification of long non-coding RNAs were already discussed elsewhere [24]. In the context of a screen conducted on genomic alignment data, it has to be noted that large parts of the genome might not be transcribed at all, or could be specific to certain cell types [25, 26], thus requiring the introduction of another classification label for non-transcribed regions. Classification of transcribed sequences is only viable if the available transcriptome data is sufficiently accurate to enable the assembly of full-length transcripts, which is not always the case. Furthermore, modeling an ambiguous background is of utmost importance when screening sequences of unknown function, as is the case with genome-wide screens for identification of potentially transcribed loci of interest and their hypothetical function.

Machine learning methods are utilized in many automatic annotation pipelines in the field of comparative genomics [27, 28]. A key issue, however, remains the inability of trained models to adapt to other species not sufficiently represented in the training data. This poses a serious problem since features like nucleotide frequencies, codon usage, and structure motifs are distributed unequally across different realms of life. Therefore, one-size-fits-all approaches typically cannot tap their full potential as they have to forego properties of high discriminative power specific only to a smaller set of related species.

As a remedy, we implemented `Svhip`, a novel software pipeline for the calibration of diverse machine learning models used in genome-wide non-coding and/or protein-coding RNA screens. The key feature of `Svhip` is the ability to train a multitude of different models automatically while giving full control over individual steps to the user. We offer the possibility to generate both models for the identification of structurally conserved non-coding RNA elements against an ambiguous background as well as for the differentiation between non-coding RNAs, protein-coding sequences and/or undefined others. Generated models can be used in the framework of `Svhip` itself or, in the case of two-class models, be exported to be used with the highly successful annotation tool `RNAz` [19, 29]. Long-term, we aim towards creating a diverse collection of models for various applications working out of the box, as well as enabling users to share their own models and parameter sets. This, by itself, ensures a large amount of reproducibility and accountability to individual experiments and genome screens and their resulting discoveries.

## 2. Materials and Methods

### 2.1. Test Data preparation

Two separate data sets were used in the conception and evaluation process of `Svhip`. For initial testing, we used a preexisting data set that was originally applied to evaluate `RNAz` [29] and `SISSIz` [30], which consists of 3 832 established and structurally conserved alignments with 2 to 6 sequences of non-coding RNAs sourced from the Rfam database [31, 32]. A wide range of vertebrate species is covered in this set. As a control, it also contains a set of random genomic locations taken from full genome alignments, which mirrors the non-coding set in length, number of sequences and dinucleotide composition.

To test the capabilities of our own genomic background simulation approach for future data set generation, we shuffled each non-coding RNA alignment on a column-wise basis using the `rnazRandomizeAln.pl` tool from the `RNAz` software framework. We also simulated alignments based on the non-coding subset with conserved dinucleotide composition and gap patterns utilizing `SISSIz`. Therefore, the data set consists of four subsets (non-coding, random genomic locations, column shuffled and `SISSIz`-simulated) with 3 832 alignments each. If not otherwise indicated, this is the data set used in evaluations.

A second data set is based on a current 27-way full genome alignment with the model organism *Drosophila melanogaster (D. mel.)* as a reference. The corresponding annotation in GTF format was obtained from FlyBase^1^ [33, 34]. To prepare full genome data for later assessment and comparison with the `RNAz` software, alignments were sliced into overlapping windows with the `rnazwindow.pl` software on a per-chromosome basis. We used windows of length 120 nucleotides and a step length of 40 nucleotides, generating an expected overlap of 80 nucleotides between neighboring windows. This rather large overlap is necessary to reduce the likelihood of falsely excluding genetic loci that display only very localized conservation signals. It should be noted that the parameters listed here are not mandatory for `Svhip`, but are considered optimal for most `RNAz` use cases. As we want to achieve a direct comparison with `RNAz` on a per alignment basis, we therefore followed its original protocol as closely as possible.

True labels of *Drosophila* alignment windows for evaluation purposes were assigned using the annotation in GTF format as follows: For each window we calculated the overlap with annotated transcripts, either coding or non-coding, as a fraction of the total window length. If the overlap exceeded 0.75, we assigned the corresponding label to the window. The cutoff at 0.75 was chosen to leave some room for disadvantageous window slices and to increase the likelihood of at least one window covering most genes with sufficient length for detection. The process was facilitated using a helper script of `Svhip, svhipAnnotate.py`, which is part of the software package.

### 2.2. Data processing

The data processing pipeline is explicitly intended for the preparation of raw alignment data for model training. It is not to be used for prediction runs, as the filtering steps implemented here may disrupt signals. `Svhip` accepts multiple sequence alignments in *Clustal* or *MAF* format as input. In the first processing step, these are grouped by the label assigned to them during initialization, either ncRNA or protein in the case of two-way classification or ncRNA, protein and other for three-way classification. We then use the `rnazselectseqs.pl` script for the initial selection of sequences. All sequences with a pairwise identity beyond a certain threshold (defaulting to 98%), are rejected. This is to ensure that redundancy in the data set is reduced to a minimum.

All alignments in the set are cut into overlapping windows of lengths 50 to 200 nucleotides. These windows then form the basis for the calculation of feature vectors for subsequent model training. Structure conservation is considered a prime indicator of functional RNA elements, windows representing the ncRNA subset of the data are filtered for this property. Alignment windows are shuffled column-wise and a tree representation of the most stable secondary structure for each sequence is calculated utilizing `RNAfold` of the `ViennaRNA package` package [35]. Subsequently, the average tree edit distance is calculated using `RNAdistance` for the window as an approximation of the remaining structure conservation after shuffling. This step is repeated for all windows and a Gaussian background distribution is fitted to the observed average edit distances. This distribution is then used to filter the input alignments by calculating an empirical *p*-value of their average tree edit distance, which is considered significant if it is lower than 0.05, see Figure 2a. This ensures that alignment fragments with a low structure conservation do not dilute the ncRNA training set, as would be the case with, for example, lncRNAs containing long unstructured stretches. In highly conserved alignments, the column-wise shuffling ensures a reduction in structure but not sequence conservation. An acceptance statistics for 100 randomly chosen alignments from the pool of our 3 832 non-coding RNA alignments or genomic locations, respectively 2.

**Figure 2:**
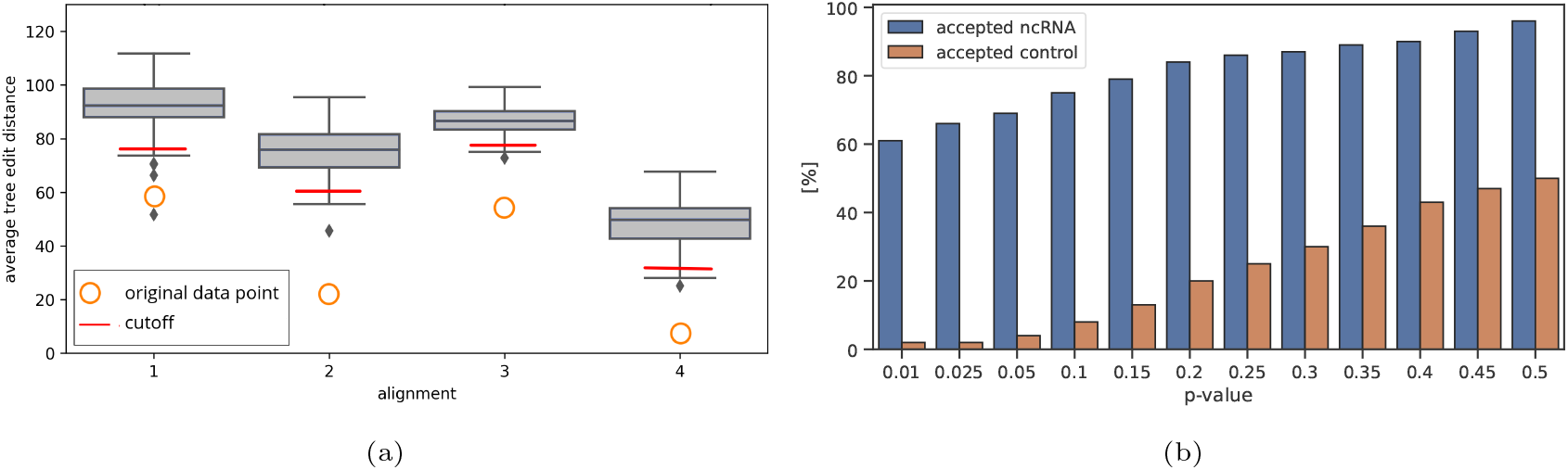
The effect of column-wise shuffling on the average tree edit distance of the sequences in an alignment. The tree edit distance between two sequences of the alignment is derived from the tree representations of the secondary structure corresponding to the minimum free energy states. Figure 2a depicts the average tree edit distances of four selected alignments (o) and the resulting distribution obtained by column-wise shuffling of these alignments (box-plot). Red lines indicate the estimated cutoff for statistical significance with a *p*-value of 0.05. Figure 2b illustrates possible outcomes of this filtering approach using different *p*-value cutoffs. The figure compares the acceptance rates of 100 randomly chosen strongly conserved non-coding RNA alignments and 100 alignments of random genomic locations as a control group. While it can be observed that the acceptance rate for conserved ncRNA elements remains relatively high at 60% even at a strict cutoff of 0.01, the acceptance of control alignments becomes increasingly unreliable with higher *p*-values. For this reason, a low *p*-value cut-off is usually preferable to reduce the false positive rate.

If no negative set representing the genomic background is provided on program initialization, a synthetic negative set will be generated based on the input data. To achieve this, `SISSIz` is applied to simulate alignments controlled for dinucleotide composition and gap pattern. Tests show that `SISSIz` generated alignments can be used as a valid approximation for randomly selected genomic background yielding comparable levels of structure conservation and minimum free energy (Figure 4). This makes it viable to simulate an arbitrary background, while mostly conserving species-specific nucleotide distributions based on the input alignments. We also investigated the viability of using column-wise shuffled versions of the input alignments as negative sets (Figure 4a). For this we used `rnazRandomizeAln.pl`, which also attempts to conserve local conservation and gap patterns during shuffling.

The alignment windows that were not filtered are then considered as representative of their respective class and serve as the basis for the generated training set. We use five features to capture essential information from these alignments while keeping redundancy at a minimum.

#### Structure conservation index

The structure conservation index (SCI) is a metric for the alignment-wide presence, size and frequency of sites that are conserved in their secondary structure. The SCI compares the MFE of individual sequences with the consensus sequence MFE. Significant deviations indicate structured elements that are not present in other sequences. Thus, a value greater or equal to 1 represents a very strong level of conservation, while a value close to 0 indicates none. The SCI for the alignment *A* is estimated by

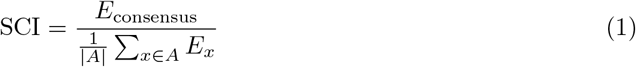

where *x* ∈ *A* are the |*A*| individual sequences in *A, E_x_* denotes the MFE of *x*, and *E*_consensus_ is the MFE of A’s consensus sequence.

#### z-score of minimum free energy

The minimum free energy is a reliable indicator for the presence of paired bases and, hence, secondary structure. A point of interest in the search for conserved non-coding RNA elements is identifying (sub-)sequences with an MFE that significantly deviates from that of sequences with equal nucleotide composition but no notable secondary structure. To detect these, the *z*-score of MFE is used. It can be calculated using one of the following two procedures. In the first case, a given sequence is 100 times dinucleotide shuffled using the Altschul– Erickson algorithm [36] and the MFE is calculated for each resulting sequence. For the resulting distribution of random MFEs, the standard deviation *σ* and mean *μ* are calculated. Finally, the *z*-score of the given MFE *X* is computed as

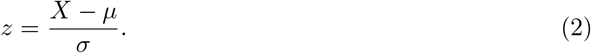

The mean normalized *z*-score of the MFE for the full alignment can thus be expressed as:

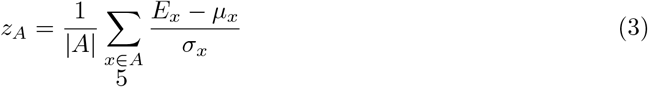

where *σ_x_* denotes the standard deviation, *E_x_* the MFE and *μ_x_* the mean for sequence *x*. This, however, is a highly expensive operation in terms of computation time per input alignment as it effectively requires the sampling and estimation of the unimodal distribution for each sequence in the alignment. For this reason, we implemented support vector regression models that estimate the expected standard deviation and mean based on fast-to-calculate, sequence-intrinsic features. These are: all 16 dinucleotide frequencies, C to G+C nucleotide fraction, A to A+U nucleotide fraction and the normalized length of the sequence. The models were trained on multiple large sets of synthetic sequences representing GC-content fractions in 0.1 intervals from 0.2 to 0.8 and sequence lengths from 50 to 200. Yielded values are within an acceptable deviation from those obtained with the exhaustive approach outlined above, see Figure 3. In our use cases, this approximation yields sufficiently accurate results. It is however possible to manually override the prediction process and enforce a shuffling and distribution sampling for all input alignments for maximum accuracy. It should be further noted that both approaches yield reasonably stable results: The SVR will naturally return identical values for identical input vectors, the mean MFE values obtained by manual sampling in our tests typically oscillated by less than 5% for 100-fold resampling (Figure S1).

**Figure 3:**
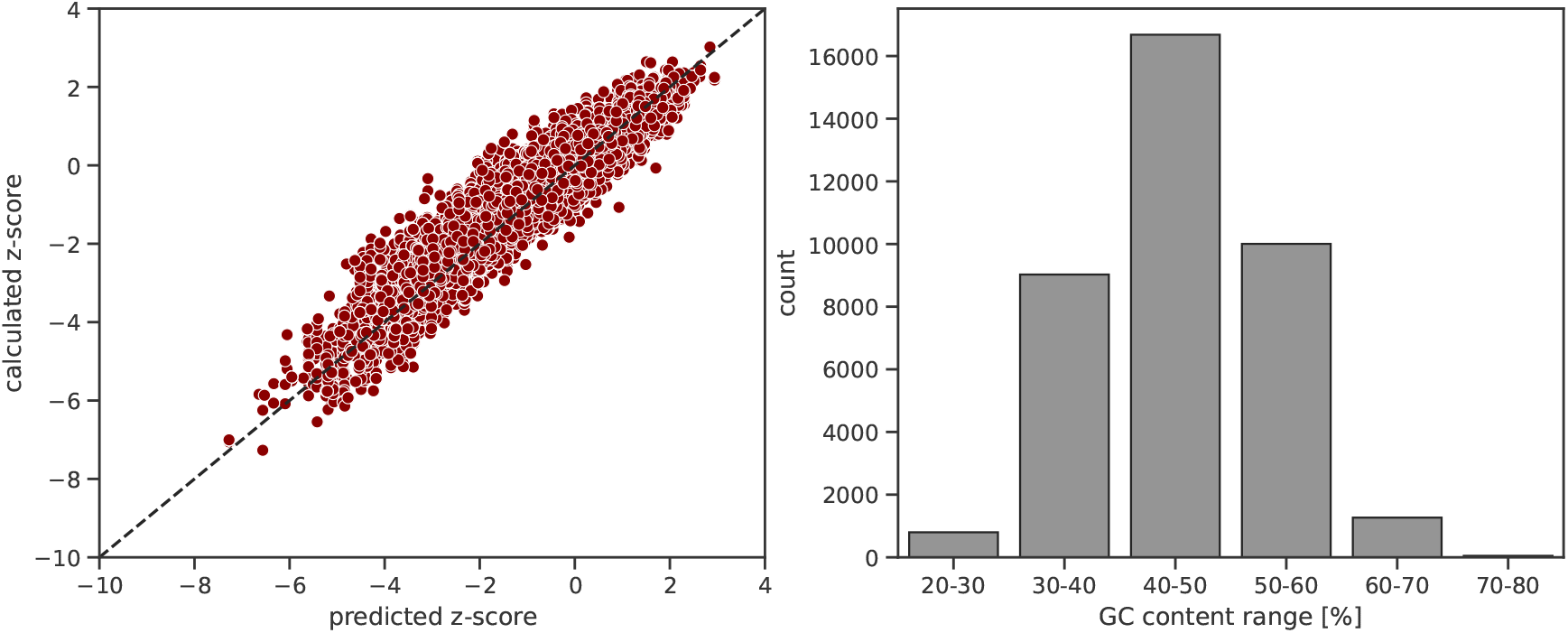
Performance of the support vector regression engine trained to predict the standard deviation and averages of minimum free energy (MFE) values of sequences with a given mono- and dinucleotide composition. A test set of 24 000 sequences was randomly selected from the non-coding and the random genomic subsets of data set one. The SVR was trained on 285 527 synthetic sequences generated to account for different GC content in 10%-intervals and different A/(A+U) and C/(C+G) ratios in 5%-steps. Left: The Figure shows the predicted and the calculated *z*-scores obtained from real distribution sampling. For a small number of sequences within the test set, the SVR significantly underestimates the average MFE of the shuffled sequence set (Figure S1). Right: The distribution of GC content ranges in the test set is shown. Sequences with a GC content of less than 30% and more than 70% are comparably rare in the test set. In our understanding, this accurately reflects the expected frequencies encountered in genomic test data, e. g. in the *D. mel.* genomic data analyzed as a case study.

**Figure 4:**
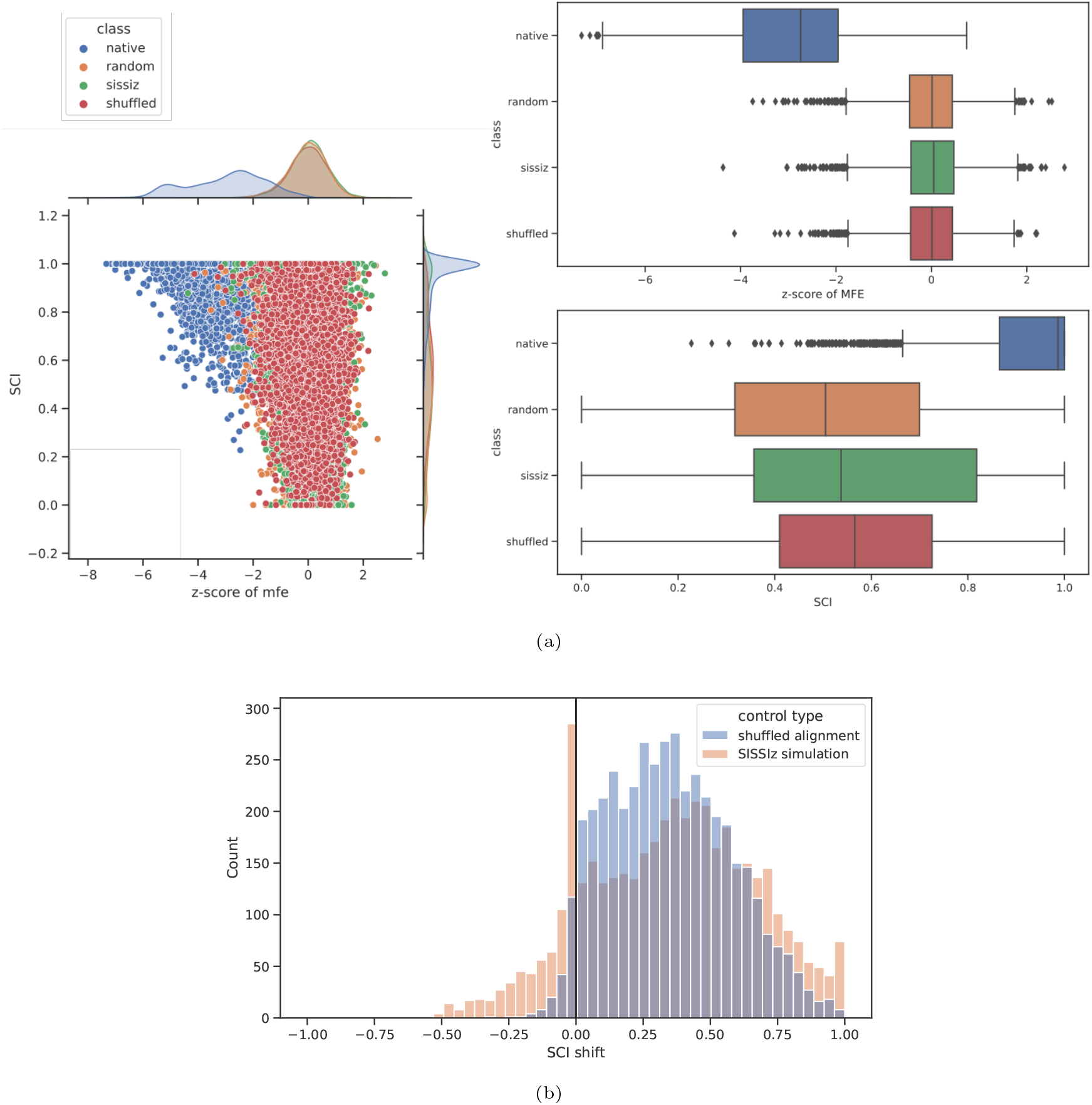
Comparison of the *z*-score of minimum free energy (MFE) between non-coding RNA alignments and different methods of sampling a negative set for the simulation of a genomic background. The *z*-score of MFE represents the statistical significance of a given alignment’s normalized free energy when compared with random sequences of equal composition. The background distribution was estimated by shuffling the sequences of the alignment 1000 times, calculating the MFE for each and fitting a Gauss distribution. The *z*-score was then calculated using its standard deviation and average. As can be seen, statistical significance is primarily achieved for the genuine non-coding RNA alignments while all three methods to generate negative sets mostly yield *z*-scores close to zero. 4b: Empirical shift in the signals of the structure conservation index (SCI) per alignment as obtained by two different methods of generating a negative set, either by shuffling of the whole alignment using the `rnazRandomizeAln.pl` script, or by simulating a corresponding alignment with `SISSIz`. For the vast majority of all alignments, both methods are reliably destroying the SCI signal. The disruptive shift in SCI is crucial for the corresponding method to be viable at simulating an ambiguous genomic background, as SCI signals are typically encountered with noncoding RNA transcripts but not with random genomic location. However, we note that alignments exist where no large offset in SCI can be achieved by simulation with `SISSIz`, see for this the high bar at an SCI shift of 0. It is likely that this is indeed caused by alignments with a very uniform composition of dinucleotides, resulting in less degrees of freedom for the generation of new alignments with similar composition.

#### Shannon entropy

The Shannon entropy *H* is a metric for the sequence variation not otherwise captured by the features outlined here. It is calculated for each alignment as

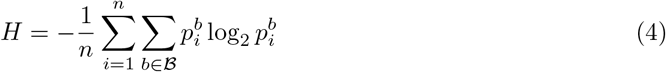

where 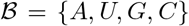 is the alphabet of nucleobase symbols, and 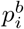 represents the empirical probability of nucleobase *b* at the i-th of (in total) *n* alignment columns.

#### Alignment-wide hexamer score

The hexamer score (Hex) is a common metric for estimating coding potential in arbitrary sequences and transcripts. A log-odds ratio is calculated based on the probability to find a given 6-mer of nucleotides in either a protein-coding or non-coding background [37, 21]. It is conceptually built on the observation that amino acids in a given protein influence their neighbors in their empirical likelihood to appear in this position. The score is largely dependent on the observed reading frame.

Our approach for the calculation of an alignment-wide hexamer score is as follows: first, the hexamer score is calculated for each sequence and reading frame using the formula

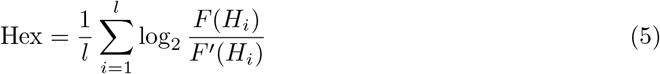

where *l* is the number of possible hexamers in the sequence, *F*(*H_i_*) is the empirical probability to find hexamer *H_i_* in a coding sequence and *F*′(*H_i_*) is the empirical probability assuming a non-coding background [21]. As these genomic backgrounds are obviously not equal for all organisms, probability models are provided for human, mouse and drosophila genomes. However, as these are only an insufficient approximation for many research cases, we also provide the script `HexBackground.py` to recalibrate these models given either an annotated genome or fasta files with coding sequences (CDS) and non-coding sequences. We then interpret the reading frame with the highest score as the “true” reading frame for the purpose of evaluation, as non-coding sequences tend to produce low scores in any frame. The average score of these maximum scores is then the alignment-wide hexamer score, Hex. We found the hexamer score to have high predictive capabilities for the purpose of identifying protein-coding sequences.

#### Codon conservation score

In terms of alignment-based coding potential estimation, the tool `RNAcode` is one of the most reliable estimators, even a decade after its initial release [38]. It relies on the simulation of a large number of potential phylogenetic trees to assess the statistical significance of the calculated score, which is computationally expensive. We built a simplified version of the underlying scoring algorithm that sacrifices some accuracy in exchange for greatly improved performance. The raw pair-wise codon conservation score (CSS) for each two aligned sequences in an alignment is then calculated as

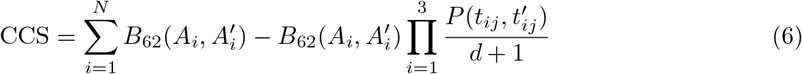

where 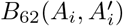 is the BLOSUM62-score of the two amino acids *A_i_* and 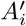 encoded by the *i*-th aligned codon in the first and the second sequence, respectively. *N* is the number of aligned codons. A rudimentary phylogenetic tree based on the input alignment is estimated using a neighbor joining approach based on pairwise sequence identity. This method was chosen for simplicity and speed, however, custom phylogenetic trees obtained with arbitrary means are supported. 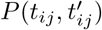 is then defined as the empirical probability to observe the mutation from the nucleotide in position *j* in triplet *t_i_* encoding *A_i_* to the corresponding nucleotide in 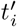 encoding 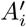 given the estimated tree. The relative evolutionary distance *d* between the sequences is estimated according to the phylogenetic tree.

The above score is calculated accounting for all possible reading frames. Faults in the alignment are skipped over by correcting for single-nucleotide deletions and frame breaks, see Figure 5. The conservation scores are stored in a matrix and used to calculate the most probable coding subregion in the alignment by calculating the maximum subarray. The score is then divided by the alignment length for normalization and used as a feature for classification.

**Figure 5:**
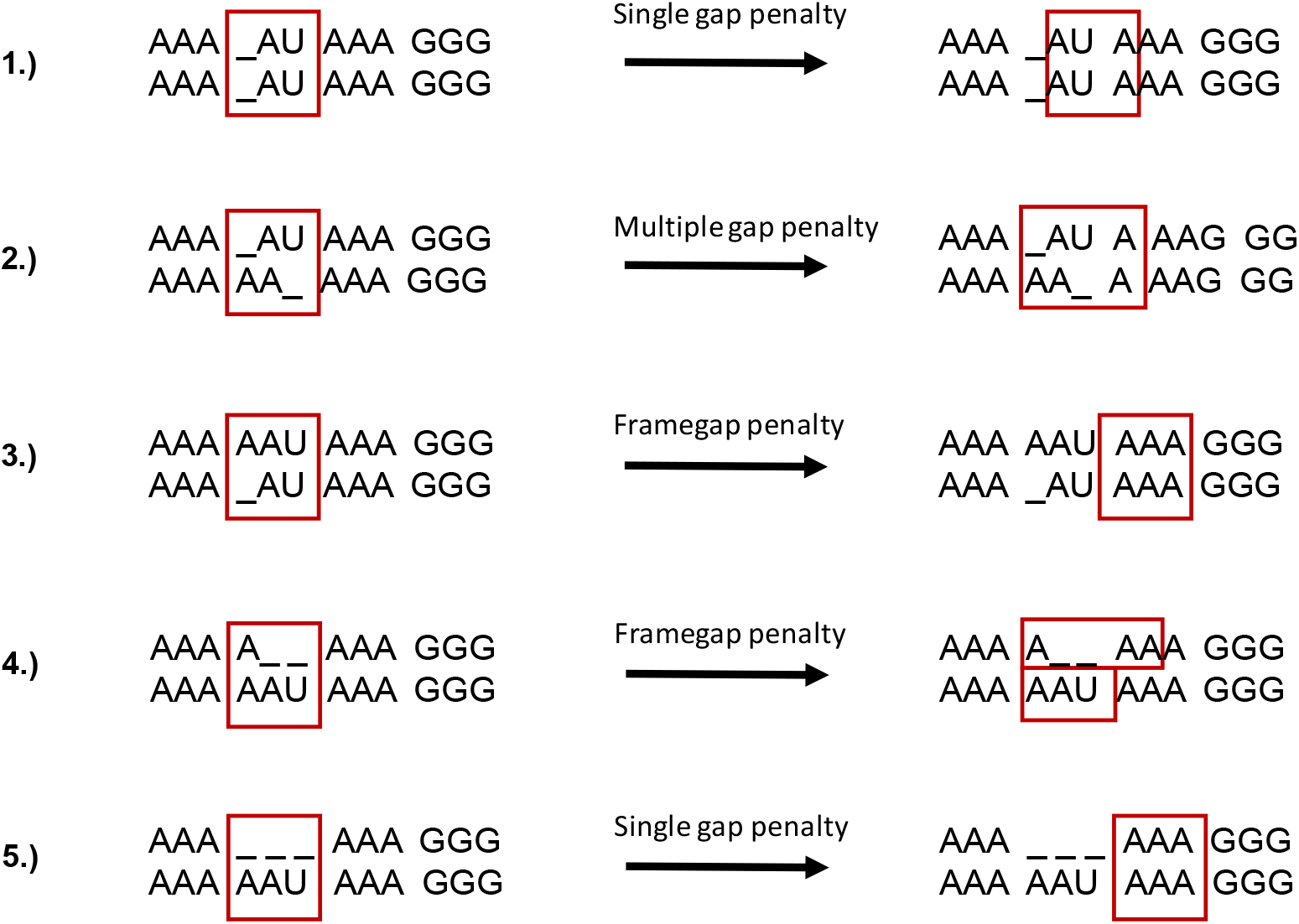
Different gap pattern handling strategies for the purpose of codon conservation calculation. The currently analyzed triplets are marked with red boxes. The left illustrates common patterns of gaps while the right hand side shows approaches how these are handled by highlighting the next analyzed triplet. 1) In the simplest case, there is just an offset by “aligned” gaps that are present in both sequences. In this case, the reading frame will simply be adjusted by the number of shared gaps and a small gap penalty will be applied. 2) If gaps occur that are not “conserved” between sequences but that sum up to the same number, the frame is corrected by being extended one-sided exactly by this number of gaps. 3) If there is a single gap in the non-reference sequence, it is likely that a sequence error is present here. In this case a constant penalty is applied and the gap is ignored, leaving the original (reference sequence) frame intact. 4) If there is one or a number indivisible by three gaps in the current reference sequence but not in the aligned sequence, we will assume the correctness of the reference sequence and the corresponding sequence frame is extended by that number. 5) If there are gaps that are multiples of three, the reading frame stays intact and is just shifted past the gaps. Only a slight penalty is applied to reflect the deletion of complete codons. Applied penalties are determined as follows: in case of a single or multiple gap penalty, the penalty has a value of −1 times the number of observed gaps. In case of a frame gap penalty, the value is fixed at −12.

The calculated features are saved in CSV format and can thus readily be accessed. Statistical information regarding feature distribution and degree of separability is provided as part of the output.

### 2.3. Model training

By default, `Svhip` supports three different types of models: support vector machines (SVMs), random forests (RFs) and logistic regression (LR). In principle, every feature set can be used to train each type of model, even though there are some qualitative differences between them. SVMs have by far the highest computational complexity, whereas LR has the lowest [39]. Furthermore, LR might not be suitable for particularly complicated classification scenarios with large overlap of feature distributions between classes, which is exemplified by the fact that many biological classification problems are not linearly separable. However, the lower training and classification time might make using an LR model worthwhile for a preliminary screen or when using an exceptionally clean data set. The question *whether SVMs or RFs* are fundamentally better in terms of raw accuracy for genomic applications is currently unanswered, even though both were capable of separating the same data sets equally well in our own studies when given appropriate training data. When we trained RF classifiers instead of SVMs using the same training data and using the same test data, we received marginally worse accuracy results (data not shown). However, we believe that the size of the data sets is too small for generalized conclusions here.

`Svhip` offers automatic hyperparameter optimization for all models using either a grid search or a random walk based approach. Both options are fully customizable in their search depth and number of cross-validation steps, however, we found that more than five cross-validation iterations typically do not offer a significant improvement in accuracy given the features used (data not shown). Generated models are saved in a binary format and can be loaded using the `predict` option. We also note that, under certain circumstances, the expected increase in accuracy by hyperparameter optimization might be marginal when compared with the quality of the training data. Parameters associated with training, including hyperparameters and scaling parameters for normalization are saved together with the model and loaded as required for prediction purposes.

### 2.4. Prediction with generated models

`Svhip` classifies alignment data by either employing one of the built-in prediction models, or by resorting to a custom model previously generated using the `features` command as described above. For the classification of input data, the alignments have to be cut into overlapping windows to achieve accurate results, analogous to their preparation for feature calculation. An alignment length of 50 to 200 nucleotides with a sliding overlap of 40 to 80 nucleotides yields reliable results for the cases presented here. For the preparation of windows, the software `rnazwindows.pl` of the `RNAz` framework was applied. The predicted class labels are assigned to the data sets on a per-window basis. Furthermore, an overall class probability is reported which, in the case of SVM classifiers, is based on a regularized maximum likelihood score [40]. RFs natively support the calculation of class probabilities as a fraction of class votes by trees in the ensemble, and LR inherently calculates probabilities by virtue of being a likelihood function fitted to binary observations.

### 2.5. Quality assessment of Svhip-trained classifiers

To assess the quality of the models generated by our software, we first constructed a classifier from curated Rfam data and compared the performance with the unmodified `RNAz` software. Sensitivity and specificity were estimated on the same data set used in the original assessment of `RNAz`’s accuracy [29] (see section 2.1). The training set was assembled by manually selecting the first 50 Rfam alignments with known and significant secondary structure as evident by the literature, see Table S2. The alignments also had to contain more than 50 sequences. If this was the case for the seed alignment, it was used directly, otherwise we chose the regular alignment instead. These were downloaded as fasta files, and a corresponding data set was generated using `Svhip` with standard parameters. Based on this data, a SVM classifier was trained, using the builtin grid search hyperparameter optimization with a range of 1 to 10 000 with a logarithmic increment for the cost parameter C of the SVM. As this parameter in general governs the weight of misclassifying any given observation in a support vector machine, it is vital to optimize it for any modified training set. The same range was used as a baseline for the gamma parameter, however the auto-scaling option as provided by the `scikit-learn` [41] package was also added to the test range and proved to be the most efficient. The auto scaling sets the gamma value to one divided by the number of features and is, in many cases, a good estimate [41].

We note that far more care could have been taken to ensure the selection of high-quality and representative input data sets. However, an important point to demonstrate here is the underlying hypothesis that the data preprocessing pipeline as implemented in `Svhip` already optimizes the data set for maximum discriminative power and lowest redundancy. For this reason, we also compared the quality of the generated classifier to one using the same input data but with most of `Svhip` preprocessing steps deactivated. This was achieved by running the `Svhip data` command with flag `-F` set to `False` and flag `--max-id` to 100. This deactivates both the tree-edit distance based filter for an insignificant structure signal and the filtering of (almost) identical sequences, respectively. The expectation here was a statistically significant reduction of the resulting capability to differentiate positive from negative class instances, presumably below both the optimized `Svhip`-generated classifier and `RNAz`. Raw classified data is contained in the supplement ^2^.

### 2.6. Full genome screen on Drosophila melanogaster

To acquire a reasonable estimate of the expected accuracy of a `Svhip`-trained classifier and showcase a basic experiment with a retrained classifier, we performed a full genome screen on *D. melanogaster* full genome alignments in Multiz format and compared predicted loci with a contemporary annotation. Alignment data for *D. melanogaster* was prepared as outlined in section 2.1. Two different `Svhip`-trained classifiers were utilized for this study: one already trained and tested in the previous section, and one with added data for protein-coding sequence differentiation.

Training data for the protein classification was assembled as follows. Coding sequences were sampled from the InsectBase [42]. To this end, we randomly selected 400 protein-coding genes from the *Drosphila* annotation and then downloaded all corresponding sequences from the database. To further reduce the amount of data, we limited ourselves to the *Ephydroidea* (containing *Drosophilids*) and *Oestroidea* superfamilies.

## 3. Results

We tested the `Svhip` pipeline and derived models for a range of different test sets and scenarios. The primary goal here was to properly estimate the specificity and expected false positive rate on both alignments of known non-coding RNAs as well as from the context of a full genome screen on *D. melanogaster*. Specifically, we also wanted to estimate the prediction quality on non-coding RNA data against the `RNAz` framework, which shares several of the implemented machine learning features, but comes with a hard-coded decision model, not allowing for retraining of the underlying classifier. In particular, we demonstrate that our preprocessing workflow for the raw training data does in fact yield high-quality positive instances, while the generation of synthetic negative data successfully reduces false discoveries to a minimum. We also show the separability of coding and non-coding data in an alignment context using our features designated for estimating coding potential.

### 3.1. Workflow and commands

`Svhip` offers different modes of action, denoted as `data, data3, training, features` and `predict`. The first two serve to generate training instances for the machine learning engine from multiple sequence alignment data, either assuming a two-way (structurally conserved noncoding RNA or protein-coding vs genomic background) or three-way classification (non-coding RNA, protein-coding and ambiguous background), respectively. The `data` and `data3` commands also evaluate the suitability of a given set of alignments for training the classifier by analyzing properties such as the average minimum free energy, levels of conservation and the overall quality of separation achievable with selected features. Every step of the pipeline is accompanied by graphical output to facilitate the assessment of the results and potential issues (Figure 6). The training command computes a classification model based on a data set generated with either of the previous commands. The remaining two modes, `features` and `predict`, serve to calculate the features required for prediction for alignment data, and to perform the actual classification using an existing model, respectively.

**Figure 6:**
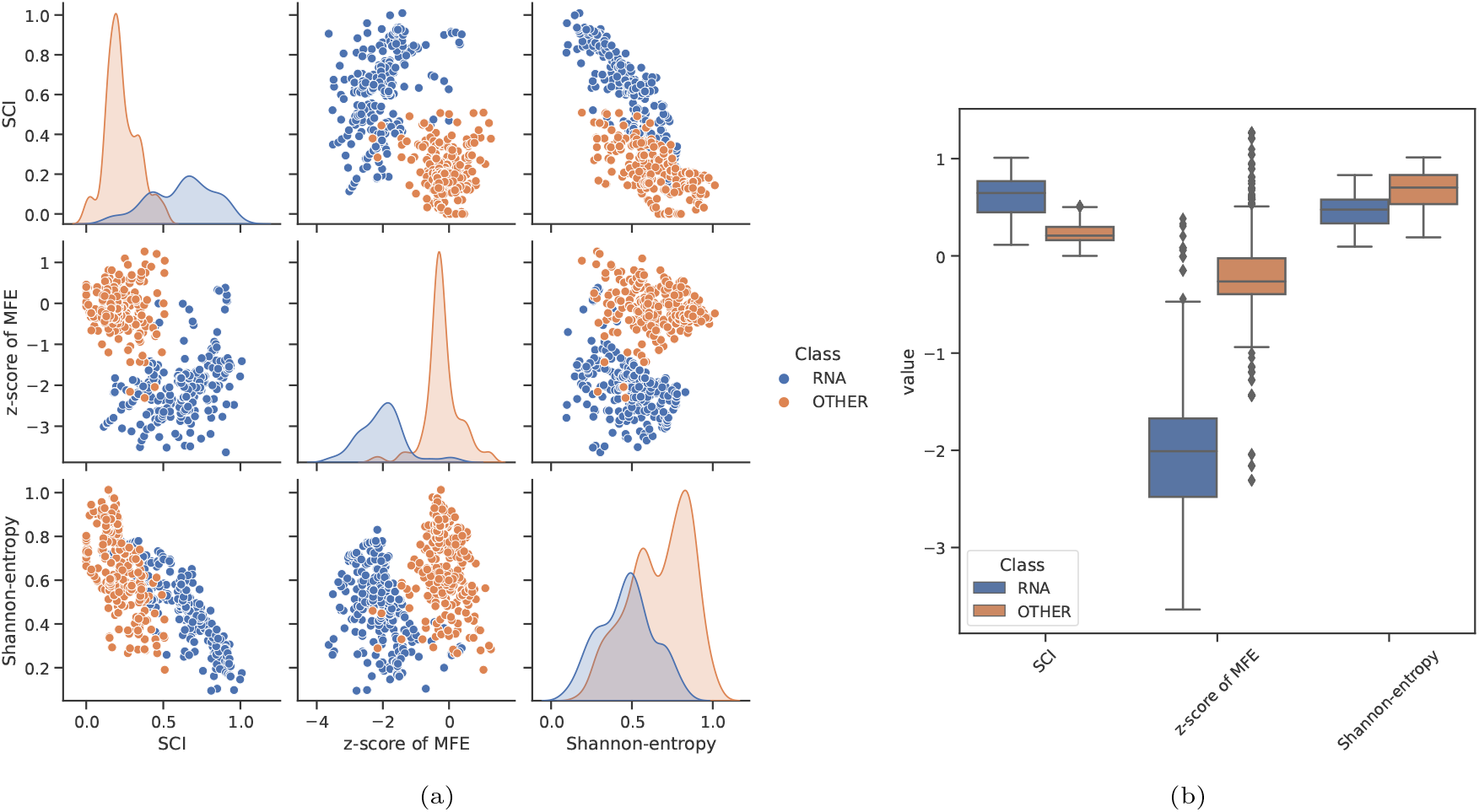
Graphical output automatically generated by `Svhip`, exemplified on an alignment of bacterial RNAse P RNA sequences (Rfam ID: RF00010). Every data point represents a generated alignment window incorporated into the training set (see supplementary Table S2). For a comparison using an input file containing both eukaryotic and prokaryotic sequences as input, refer to supplementary Figure S4. The graphical reports serve as a first indicator for the assessment of input data quality and separability of relevant features, thus hexamer score and codon conservation distributions are not shown for the RNA example here. Figure 6a: Scatter matrix showing relations between individual features as well as density plots. As can be observed, a good separation between native non-coding sequences and the `SISSIz`-generated control set can be achieved based on SCI and the *z*-score of MFE. The discrepancies between class cardinalities, as inferred from the density plots in the diagonal, can be attributed to the structural conservation filter employed in the pipeline, causing a reduction of viable data points for the non-coding RNA set. Figure 6b: Box plots illustrating the distribution of individual features, allowing the same observations in a more streamlined way. These reports may also serve as a basic exploratory analysis of newly assembled alignments, taking note of statistical significance of secondary structure properties as compared to the automatically generated control set.

### 3.2. Protein-coding features

We generalized the hexamer score, commonly used to estimate coding potential of individual sequences, to be applicable to alignments. Furthermore, a reduced version of the scoring scheme applied in `RNAcode` is implemented in the codon conservation score that sacrifices some accuracy for substantially improved speed, see 2.2 Data processing and Figure S2 for a comparison of run times.

As the codon conservation score was derived as a simplified version of the algorithm employed by the `RNAcode` software, we did a comparative analysis between both approaches on the *Drosophila* genome alignments (see section 2.1). All alignment windows containing at least three sequences were screened using `RNAcode` and their respective codon conservation scores were calculated. The restriction of the sequence number was necessary as `RNAcode` does not natively support the calculation of scores for alignments of only two sequences. As `RNAcode` reports scores for multiple possible subalignments while the codon conservation score is alignment-wide, calculating only the sum of the maximum subarray, only the best `RNAcode` hit was taken into account for this comparison. The resulting Pearson and Spearman correlation coefficients of 0.49 and 0.5 indicated only a weak correlation of both scores. In part, this can be explained by the fact that the codon conservation score takes the full alignment length into account and not just the length of the subsequence with the highest individual score. A more relevant question, however, is whether both scores are suited metrics to tell coding and non-coding sequence alignments apart. Using annotated alignment windows as a basis and separating them into *coding alignments* and *other alignments,* we performed a Wilcoxon rank sum test on both sets of data and metrics, respectively. Using this approach, the null hypotheses that both sets of measurements originate from the same underlying distribution can be rejected for both metrics with a very high confidence as indicated by *p*-values smaller than 10^-15^. Thus, while there are certain intrinsic differences in both approaches, they do achieve a strong differentiation between coding and non-coding sequence alignments.

The approach utilized by `RNAcode` also yields higher raw discrimination power in comparison to the hexamer score (data not shown) when screening explicitly for the highest scoring, connected area in an alignment. In this sense, a protein-coding training set that was pre-screened using the `RNAcode` approach would, in the most cases, be of higher reliability than one relying exclusively on the alignment-wide hexamer score. The former is however associated with significantly higher computational cost in an already computationally expensive pipeline. We therefore leave the decision for either of the methods to the user.

By combining information of InsectBase and FlyBase, we generated protein-coding alignments and investigated the distribution of the alignment-wide hexamer score and codon conservation score in a coding and non-coding context. The codon conservation score shows a very strong separability of classes just based on this feature alone, while the alignment-wide hexamer score exhibits a moderate performance, Figure 7. The separation is much stronger in the training data set than for the genomic alignment data. This, however, is to be expected as a large number of genomic alignment windows naturally overlap with non-coding sequences, thus reducing the overall yield. The training instances can therefore be seen as an ideal classification set, depending on input data. It should also be kept in mind that both features measure different properties of coding sequences and thus naturally might produce results of varying specificity depending on the input data. In particular, the hexamer score measures the relationship of codons and their immediate neighbors, while the codon conservation score estimates the overall retention of or selection for coding potential in an evolutionary context. It has to be noted that, in this classification task, no single feature carries the full information for a correct label assignment. In the most cases, all of them have to be considered in the context of the remaining parameters.

**Figure 7:**
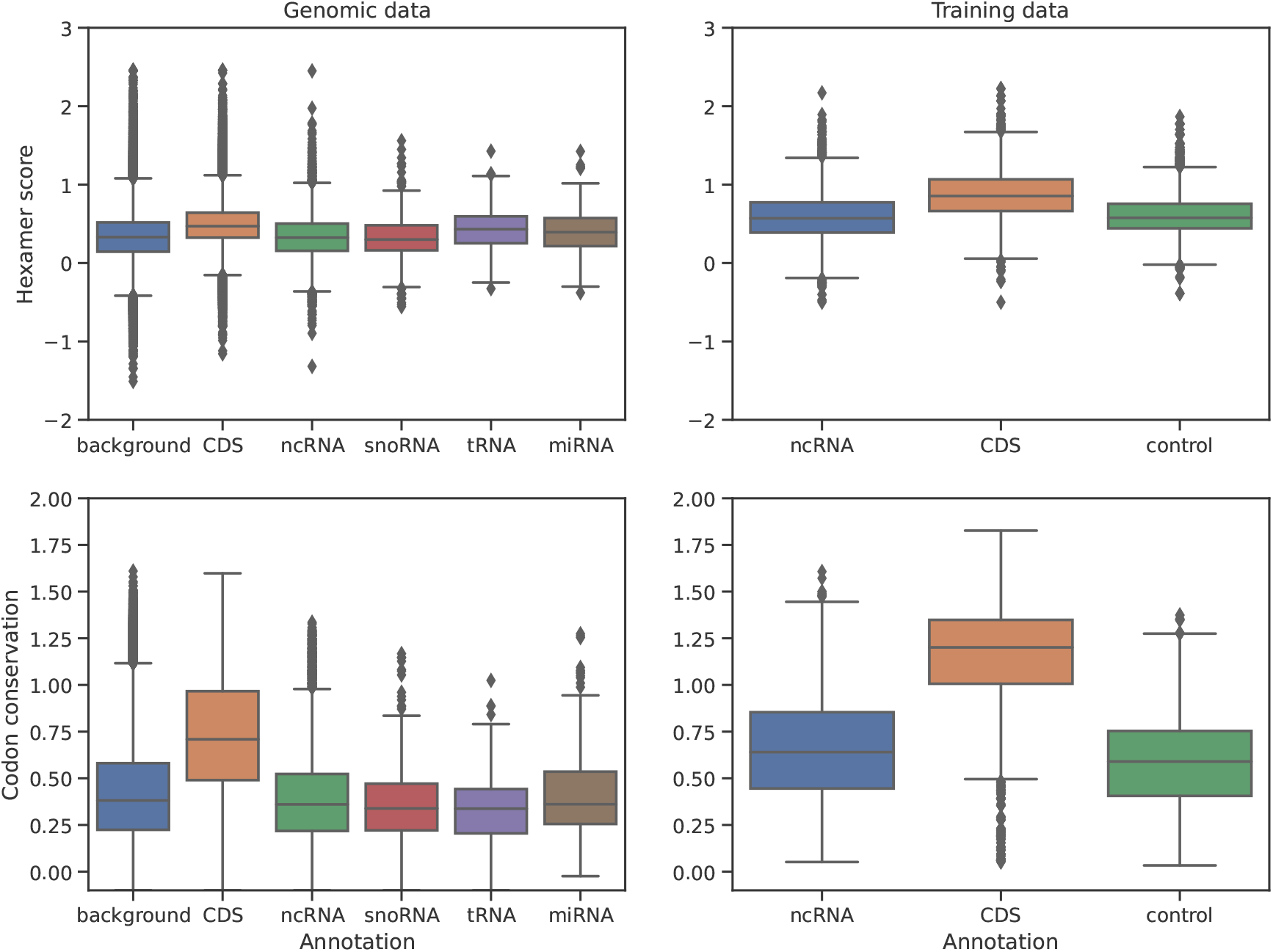
Distribution of the alignment-wide hexamer score and the reduced codon conservation score across different gene classifications and the genomic background. Classification performance on *D. melanogaster* Multiz genome alignments (left) and applied to protein-coding alignments obtained from the InsectBase, see 2.2 Data processing, when compared with the ncRNA classification training set (right). In both cases, the codon conservation score is significantly higher in most sequences annotated as a known coding sequence. To a lesser extent, the same is true for the hexamer score.

### 3.3. Analysis of two-way classification performance on alignments of known ncRNAs

Accuracy, sensitivity and specificity of `RNAz` and two `Svhip`-trained classifiers were compared on the original `RNAz` test set, the difference between the latter two lying in their optimization by filtering for secondary structure significance of the input set. The optimized classifier in this case refers to the classifier trained applying this filter method and serves to illustrate the relative effectiveness in increasing the specificity by comparison (see section 2.5). All three classifiers show a high degree of differentiation between true positive and true negative instances. Generated ROC-curves further support this impression as all three classifiers cover an area under the curve of 0.98, see Table 1. However, the ROC metric compares only relative label probabilities assigned to the test instances, and it should be understood as supplementary to the raw assigned labels, see Table S1. In terms of false positive rate, the optimized `Svhip` classifier surpasses both the `RNAz` and the non-optimized classifier by a notable margin, see Table 1 and Figure 8.

**Figure 8:**
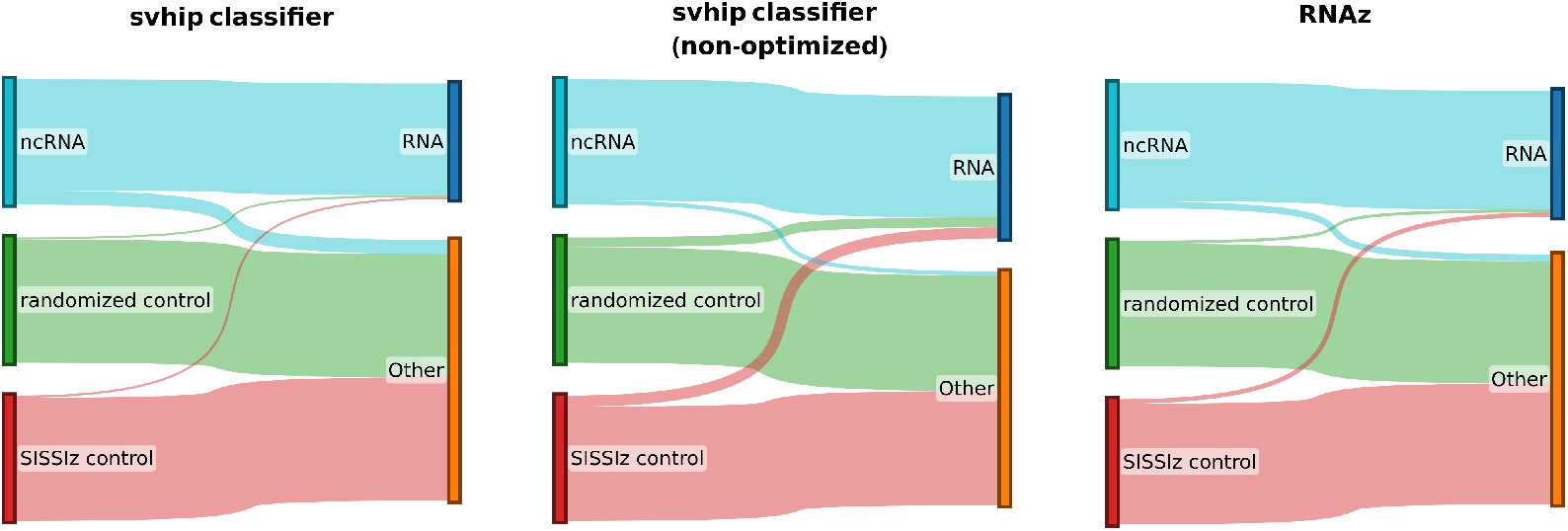
Label assignment of the three classifiers: A optimized `Svhip` classifier and another `Svhip`-trained classifier using the same training set but with no structure conservation significance filter, respectively, and RNAz. The test set contained 3 832 alignments of known ncRNAs and two control sets of (i) alignments simulated with `SISSIz` and (ii) column-wise shuffled alignments. Both control sets were based on the native ncRNA alignments. The raw numbers of classifier assigned labels is summarized in Table S1.

**Table 1:** Evaluation of classifier performance using two `Svhip`-generated classifiers and `RNAz` as gold standard. We calculated sensitivity, specificity, false positive rates, raw accuracy and the area under the curve (AUC) directly from the classification results. Note that these metrics refer to both control sets combined. In any case, the resulting numbers are very similar. As an immediate observation, the data preprocessing optimization implemented by `Svhip` notably causes a trade-off between sensitivity and specificity towards a lower false positive rate. The best value in each category is highlighted in bold.

| classifier | Sensitivity | Specificity | False positive rate | Precision | Accuracy [%] | AUC |
| --- | --- | --- | --- | --- | --- | --- |
| Svhip optimized | 0.89 | **0.98** | **0.02** | **98.18** | 95.18 | **0.98** |
| Svhip non-optimized | **0.97** | 0.92 | 0.08 | 91.98 | 93.25 | **0.98** |
| RNAz | 0.95 | 0.97 | 0.03 | 96.92 | **96.23** | **0.98** |

Based on the false positive rate on both control sets, the optimized `Svhip` classifier shows higher specificity than both `RNAz` and the non-optimized classifier, well within the lines of expectation. We note that the non-optimized classifier falls short in terms of specificity when compared to the other classifiers and that the higher sensitivity can thus be partially attributed to this. This as well is to be expected as we suggest a strong link between the selection of statistically significant training examples and overall performance. We note however that, in the context of genome wide screens, a low FPR is in many cases more valuable than a high sensitivity. This is the case for two reasons: Firstly, using a sliding window approach, the average screened genome will be separated into millions of overlapping windows, leading to an expected false positive classification of thousands of windows. Considering this, even a reduction of FPR by one percent can be considered beneficial, as the experimental validation of each individual discovered locus is substantially more costly than the genome screen itself. Secondly, as most genomic loci are larger than one window of around 120 nt, there is a reasonable chance for at least strongly conserved subsequences of the full locus to be classified correctly, such that many candidates can be identified even at lower sensitivity. For these reasons, we strongly suggest an optimization towards the lowest possible FPR feasible for a given use case. Aside from these considerations, we also note again that the analyzed classifiers are not optimized in terms of initial input data and serve more to showcase the overall capabilities of the pipeline. Alternative avenues of investigation here include for example the specific selection of only certain types of non-coding RNA as input, or a restriction to certain species of origin.

### 3.4. Analysis of two-way classification performance on Drosophila genome screen

Given the above observations, we decided to use only the optimized `Svhip` classifier for a non-coding RNA screen on the genome of *D. melanogaster*. We also expected that the overall classification accuracy of structurally conserved non-coding RNA elements would further increase when also considering protein-coding sequences as a third class. This approach, however, cannot be compared to `RNAz` directly, which exclusively screens for conserved RNA structures in an otherwise ambiguous genomic background and does not take protein coding sequences as a separate class into account. This latter fact makes a second comparison necessary, with `RNAz` on the one side, the better-performing, `Svhip`-generated classifier on the other, and a third classifier that differentiates non-coding RNA, protein-coding and background genetic locations, which will be discussed in the following section.

We performed genome-wide screens based on Multiz alignments with *D. melanogaster* as reference and the current annotation from FlyBase v2 as ground truth [34, 33]. When comparing total genes covered by preprocessed alignment windows with those recovered by individual classifiers, we note that the `Svhip` classifier (described in the previous section) performs equally well or even outperforms the `RNAz` classifier in recovering snoRNAs, miRNAs and tRNAs, Table 2. This demonstrates that the `Svhip` pipeline is well capable of isolating high-quality training instances even from suboptimal input data. It has to be noted, however, that it is difficult to correctly determine the false positive rate of the classifiers, especially for the scenario of a full genome screen. This is primarily caused by the lack of gold standard data that would guarantee correctness of the used background annotation, and the higher abundance of spurious alignments containing non-homologous or very dissimilar sequences thus introducing the possibility for reduced conservation signals despite there being a genuine conserved transcript encoded here. The ambiguity and likely incompleteness of available annotations is our primary motivation for restricting the analysis to the well-studied cases of tRNA, snoRNA and miRNA.

**Table 2:**
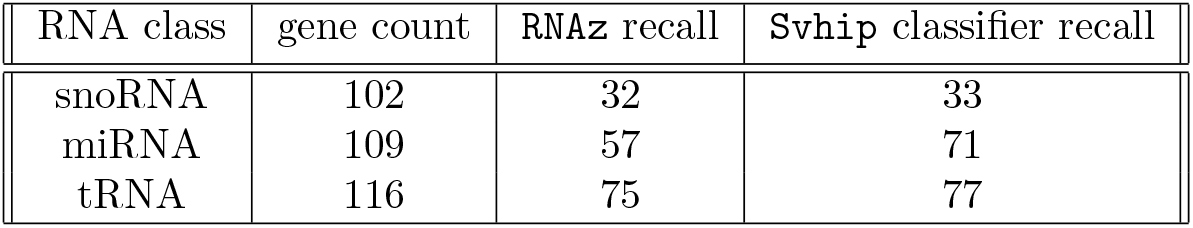
Comparison of the gene loci recall using both `RNAz` and the `Svhip`-trained classifier. Gene count refers to the number of annotated genes actually covered by sliced alignment windows, i. e. those that are actually recoverable using this strategy.

| RNA class | gene count | RNAz recall | Svhip classifier recall |
| --- | --- | --- | --- |
| snoRNA | 102 | 32 | 33 |
| miRNA | 109 | 57 | 71 |
| tRNA | 116 | 75 | 77 |

Given all generated windows, `Svhip` yielded 3 714 hits with a high confidence of at least 90%, while `RNAz` yielded 4 719. Of these, 3 483 and 3 089 are neither annotated as non-coding RNA nor protein-coding gene, respectively. If we consider the 1 128 high-confidence hits shared by both classifiers and the relative proportions of total high-confidence hits, we come to the conclusion that the false positive rate of the `Svhip` classifier is likely at least as good as that of `RNAz` for this confidence level. Together with the observations from the previous section we conclude that our classifier is about as efficient for the purpose of *Drosophila* non-coding RNA genome annotation as the `RNAz` framework.

### 3.5. Analysis of protein classification performance on Drosophila genome screen

A new classifier for the differentiation of protein-coding sequences and genomic background was trained based on the data obtained from InsectBase. The entire pool of training data from the ncRNA classification experiment, i. e. both the exemplary Rfam non-coding RNA alignments and the automatically generated negative set, was reused as a negative set. The goal of this experiment was the identification of coding regions in a full genome screen under the exclusion of everything else. While generation of a three-way classifier differentiating between ncRNA, coding sequences and genomic background is in principal possible with Svhip, it showed subpar results in this particular use case when compared with the usage of two individual classifiers (data not shown).

Of the 17835 protein-coding sequences covered by the pre-processed alignments, 13 129 were recovered by the protein classifier (recall 73.8%). Note that one gene can consist of multiple coding sequences and that such a region was counted as recovered if multiple (≥ 2), consecutive hits occurred on the same strand within that gene’s boundaries. If we only count regions as recovered if at least 50% of their length is covered by hits, we still obtain a recall of 66.1%, corresponding to 11 787 recovered regions. Obviously, these numbers are heavily dependent on the quality and quantity of the available training data and thus the resulting classifier.

Since the annotation of protein-coding genes is more mature as compared to that of noncoding RNA, the estimate of the expected false positive rate (FPR) can be considered to be much more reliable in this case. Of 477 593 alignment windows not annotated as protein-coding, 54 309 (11.3%) were classified as protein-coding loci. Assuming that at least some coding sequences in the *Drosophila* genome are not annotated, the actual false positive rate may be even lower.

Usually, protein-coding genes are substantially longer than the non-coding RNA genes discovered with this approach. In fact, there is not a single region in the *Drosophila* genome annotated as protein-coding that is not split over two or more consecutive alignment windows. This means that, in the case of protein-coding screens, isolated windows reported as hits are most certainly false positives and can be ignored in almost all cases. Applying this correction as a post-processing step reduces the false positive rate to 34 156 falsely classified windows (7.2%).

### 3.6. Chaining of classifiers for reduced FPR

In an attempt to further reduce the expected false positive rate of the non-coding RNA classifier, we investigated ways to combine the previously described classifiers. The goal here was to effectively remove as many coding sequences out of the non-coding RNA classification pool as possible as these form the largest chunk of all annotated data. It can be shown that there is a substantial overlap between potential for structure conservation in non-coding RNA and coding RNA since the conservation of base triplets tends to indirectly conserve (possibly random) secondary structures, Figure S3. In this interpretation, the potential for conservation of secondary structure is highest in genuine non-coding RNAs and lowest in the genomic background, while coding sequences exhibit an intermediate conservation level. Thus, their removal by a preceding classification step based on coding potential could substantially reduce the margin of errors in a subsequent, conservation-based non-coding RNA screening. In general, the non-coding RNA classifier outlined in 3.4 can be assumed to have a worst-case FPR of

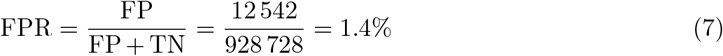

where FP (false positives) refers to non-coding RNA classification hits on windows not overlapping with a non-coding RNA annotation and TN (true negatives) refers to hits where coding sequence or genomic background windows are correctly identified as not being non-coding RNA. We explicitly counted here all potentially incorrect hits—and not only the high-confidence hits—as we assume that the probability of overlapping with genuine but not annotated non-coding RNA is higher. If the goal is to estimate an upper bound on the FPR given the classifier at hand, then all hits irrespective of assigned probability should be counted.

There are two obvious options to attempt an exclusion of coding sequences from the false positive set during classification. One is to first make a full genome screen using the protein model from 3.5, and subsequently classify the remaining alignment windows. In principal this introduces the possibility of misclassifying non-coding RNAs as coding, which then subsequently disappear from the pool of true positive non-coding RNA hits. During our analysis, however, this effect appeared to have no relevant impact on non-coding RNA classification. Unfortunately, this approach resulted only in a marginal reduction of FPR, which fell to 1.3% due to 206 alignment windows being removed from the false positive set.

The second option involves retraining the classifier after also including the protein-coding training set as part of the negative set for non-coding RNA classification, thus attempting to generate a classifier that more directly excludes coding sequences instead of only being based on the original negative set of synthetic alignments. Marking all training examples in the protein classifier training set as background (relative to the class of non-coding RNA) and retraining the non-coding RNA classifier with `Svhip` using the same parameters, yields a drastically improved FPR on the annotated *Drosophila* genome:

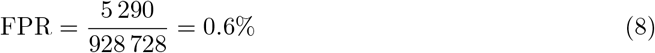

It should be noted that this classifier, as outlined in Section 3.3, only uses the features of SCI, *z*-score of MFE, and entropy, as it was noted that the more coding-sequence specific features do not substantially add to the non-coding RNA classification performance. These columns were thus excluded from the training set for this experiment.

We conclude that, depending on the use case, it may be worthwhile to manually add known negative examples to the training set instead of relying exclusively on the artificial generation of negative training instances.

### 3.7. Performance of a three-way classifier

The `Svhip` pipeline allows for the generation of a three-way classification model. We compared the efficiency with the chaining of multiple classifiers for different purposes as described in the previous section. For this experiment, the same training data was used as for training of individual classifiers. We note, however, that the three-way classifier performs notably worse using the same training data in a direct comparison. For instance, the number of correctly recalled protein-coding regions drops from 13 129 to 9 798 using this approach and the FPR becomes 5%. Due to these notable drawbacks, we limit the more in-depth analysis to the previously presented approach. On the other hand, preliminary attempts on yet incomplete training data yielded promising results for some more specific applications. We are therefore convinced that this approach may be a viable tool to consider for certain use cases. A more detailed analysis, however, is beyond the scope of this initial report.

## 4. Discussion

The `Svhip` software was used in different classification scenarios to illustrate the applicability with different research goals in mind. In principal, it can serve as both a tool for a quick screen of potential coding regions as well as for an in-depth analysis of non-coding RNAs across a variety of species. It can be demonstrated that it is also applicable in a more specialized scenario such as the search for high-confidence hits for coding potential in genes already assumed to encode non-coding RNAs with different biological functions. To our knowledge, no other tool currently unites these functionalities and such an investigation would typically require to make a screen with one specialized tool and another, already established annotation in mind. This use case has substantial downstream potential, as there is evidence that non-coding RNAs containing proteincoding subsequences are yet not close to being fully investigated [43, 44].

Our results clearly show that `Svhip` generates decently accurate classifiers with relatively high recall and moderately low false positive rates even from subpar training data. It has to be noted that most intended use cases target full genome screens instead of the classification of single alignments of full sequences, thus already making the classification task substantially harder. In cross-validation scenarios using subdivisions of our training data, we obtain recalls and false positive rates that are still markedly superior to the data obtained from a genome screen, as Sections 3.3 and 3.4 indicate. One problem here is the slicing of genomic regions into overlapping windows, a process that inherently destroys certain conservation and structure signals at the window boundaries. An important, yet insufficiently addressed problem is the classification of long non-coding RNAs in full genome screens. Long non-coding RNAs are in many ways more difficult to classify by window-based approaches because focussing on local features like secondary structure conservation becomes less viable for longer candidate sequences. Approaching this problem will be a goal of future investigation. Potential solutions include better methods to join related, but not directly adjacent hits within the same gene and improving training set composition for this specialized problem.

It must also be acknowledged that the potential and usability of the software could be further improved by continuing the development of clustering approaches for given groups of hits. As noted above, in many cases it may be insufficient to simply group adjacent hits together and declare them to be part of the same gene or region. This simplified approach is especially fallible if either a very long coding sequence or a long non-coding RNA is to be analyzed. In the former case, the large number of successive alignment windows involved naturally increases the chance for false-negative hits, and any false negative would lead to an interruption of the chain and thus to a misclassification as two protein-coding regions instead of one. In the latter case, structure conservation or the overall presence of stable secondary structure is usually not evenly distributed, again leading to accidental subdivision of loci. Both of these problems have been observed in our case studies. The situation is complicated further by the presence of introns and alternative splicing events. These shortcomings emphasize the necessity to devise better methods for identifying clusters of hits, which will be the subject of future work.

## 5. Conclusion

The software `Svhip` serves as a framework to train, test and apply machine learning classifiers in full genome screens for the annotation of protein-coding or non-coding RNA genes. We demonstrated the usage of the pipeline in a number of different scenarios and obtained comparable classification results to the `RNAz` framework when comparing non-coding RNA classification. The primary advantage of `Svhip` is the ability to in- or exclude different data sources on the fly, and to fine-tune parameters for different use cases. Further work is required to evaluate the applicability in different scenarios. One prime target is an in-depth investigation of plant genomes, many of which are currently lacking in terms of accurate and complete annotations. A particular challenge there is the high presence of gene duplication and transfer events, increasing the difficulty of the data preprocessing, and a highly volatile alignment quality and structure. We envision that the adaptability of the pipeline will serve to further improve the annotation of many different species.

## Acknowledgements

This work was partially supported by the University Leipzig PreDoc Award ^3^. The rest of the work was supported by the ScaDS.AI Dresden-Leipzig Center for Scalable Data Analytics and Artificial Intelligence, funded by the Federal Ministry of Education and Research and the Free State of Saxony.

## 6. Supplementary Material

**Table S1:**
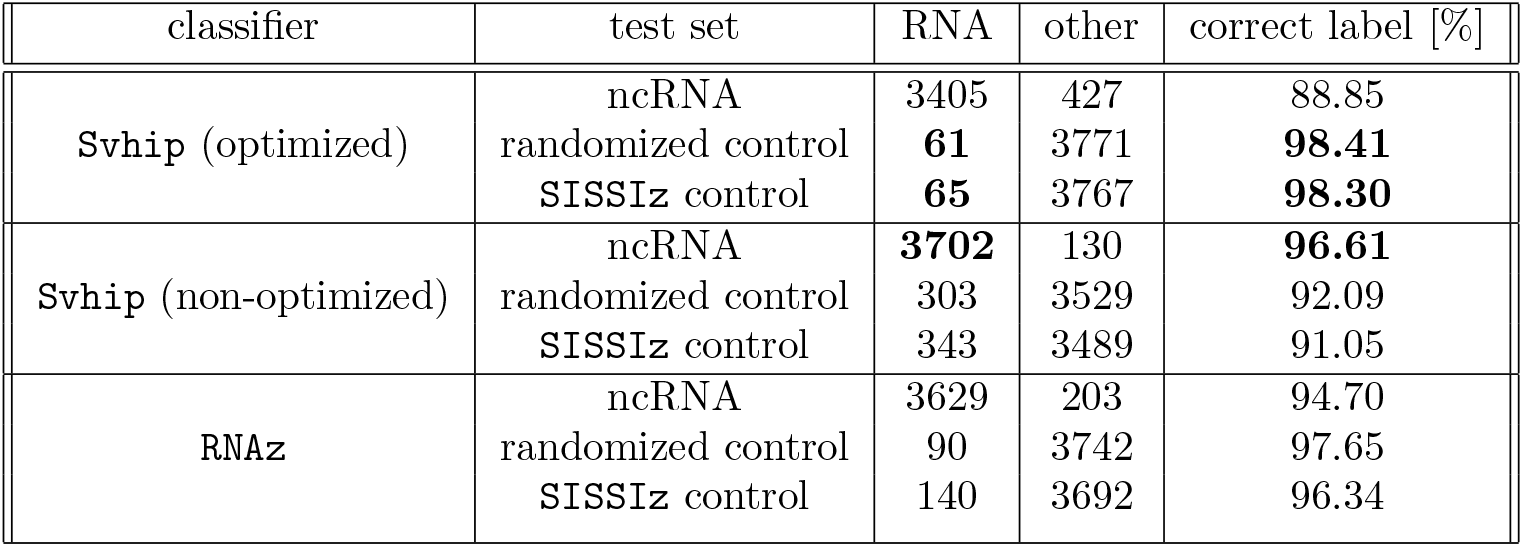
Non-coding RNA classification results comparing performance of two `Svhip`-generated classifiers using `RNAz` as a gold standard. The test set used was originally utilized in the evaluation of the `RNAz` software [29] and contains both well-established structured non-coding RNA alignments as well as alignments taken from random genomic locations as a control (randomized control in the table). We extended the control set with a set of alignments simulated using `SISSIz` to account for the fact that simulated alignments were used during training set generation (`SISSIz` control in the table). Class labels assigned were either *RNA* for supposed ncRNA alignments, or *other* for everything else. The best value in each test set was highlighted in bold.

**Table S2:**
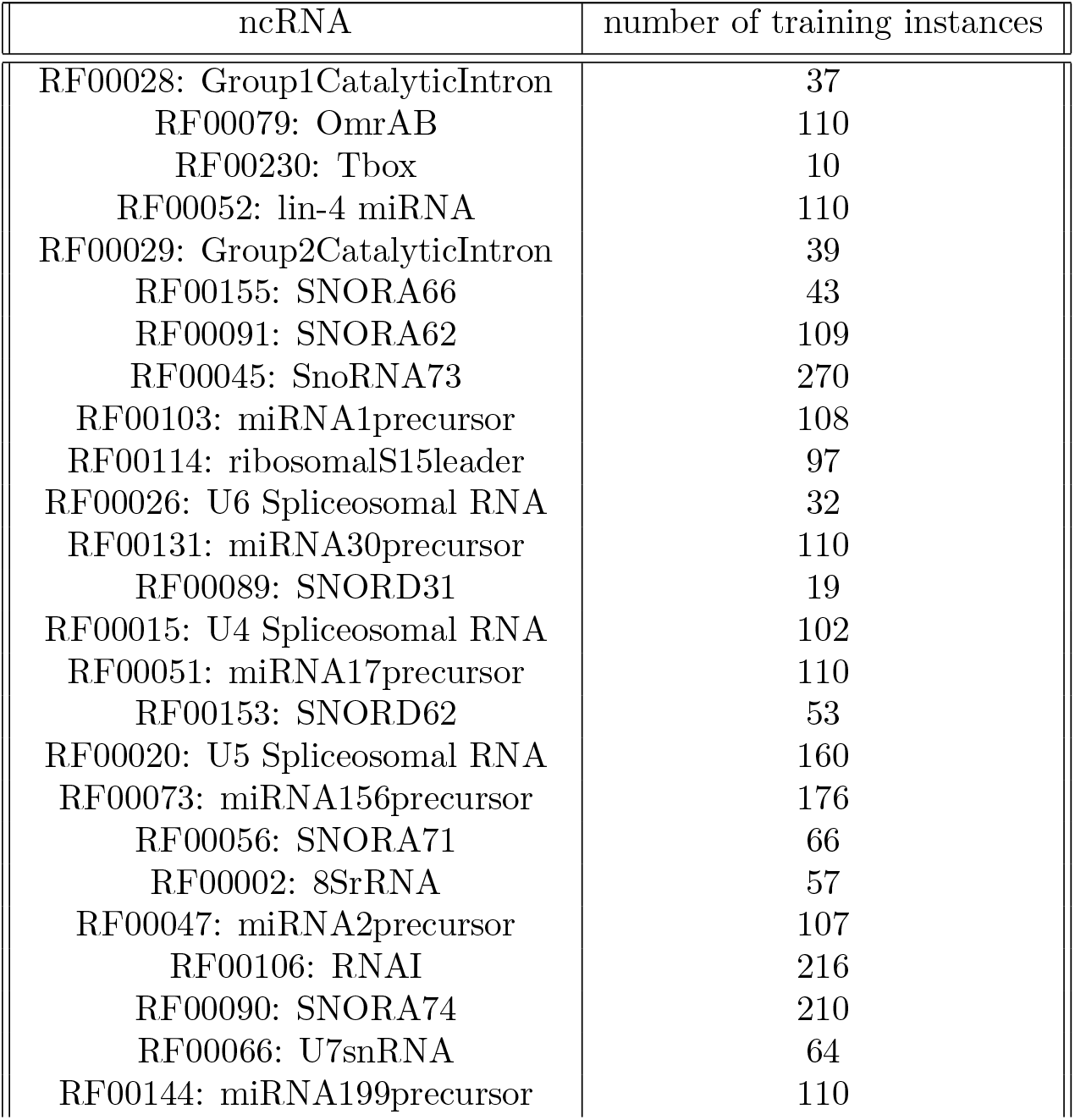
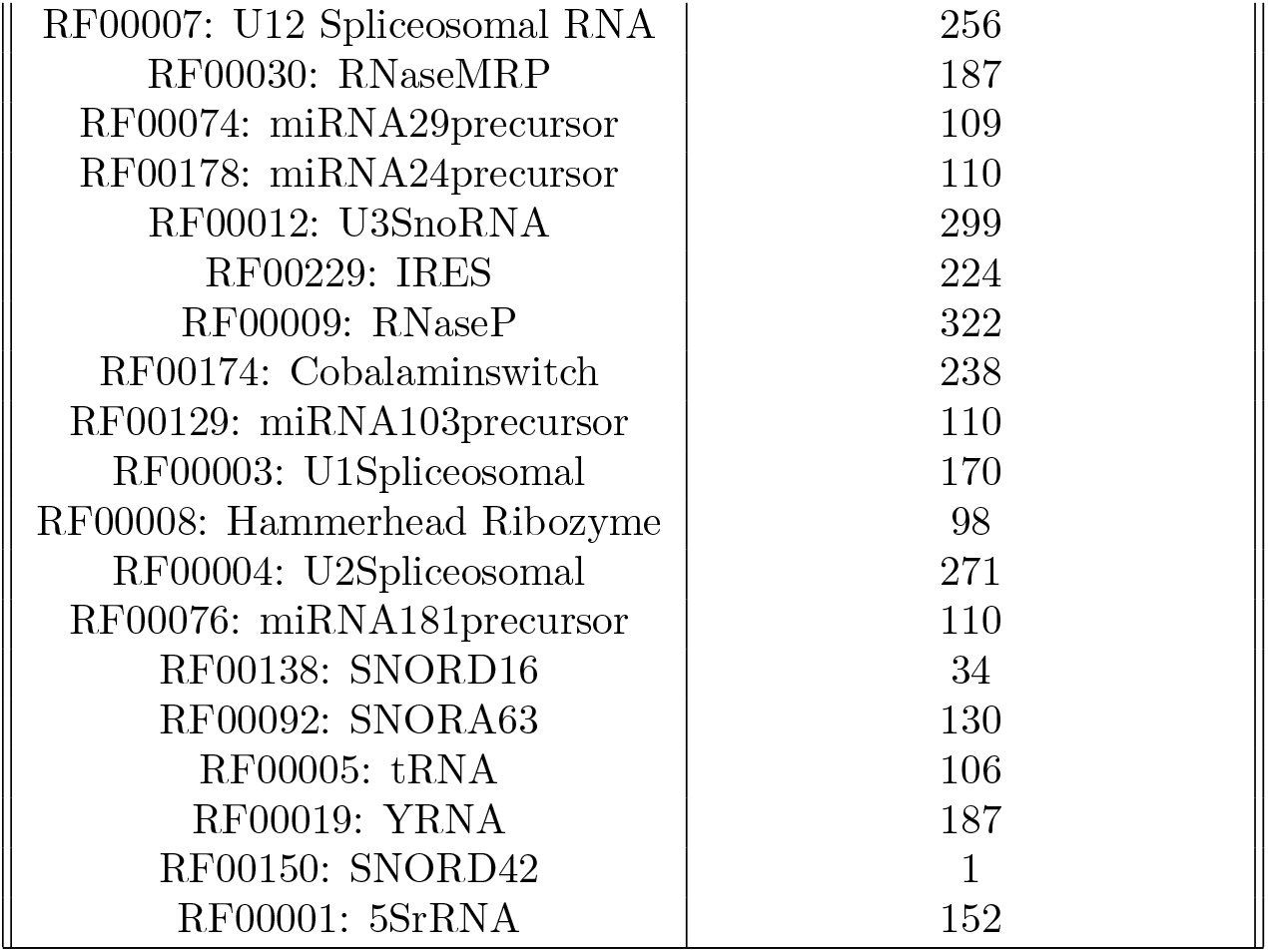
Selected non-coding RNAs for usage in classifier training as obtained from the Rfam database. We only selected non-coding RNA families that (1) show sufficient evidence of structure conservation, and (2) comprise at least 50 sequences, either in the seed alignment, or otherwise in the regular alignment. Note that the number of training instances varies with the initial number of sequences, length of alignment and number of windows reduced during filtering. RNA families that did not yield any training instances after all filtering steps were excluded from this list.

| ncRNA | number of training instances |
| --- | --- |
| RF00028: Group1CatalyticIntron | 37 |
| RF00079: OmrAB | 110 |
| RF00230: Tbox | 10 |
| RF00052: lin-4 miRNA | 110 |
| RF00029: Group2CatalyticIntron | 39 |
| RF00155: SNORA66 | 43 |
| RF00091: SNORA62 | 109 |
| RF00045: SnoRNA73 | 270 |
| RF00103: miRNA1precursor | 108 |
| RF00114: ribosomalS15leader | 97 |
| RF00026: U6 Spliceosomal RNA | 32 |
| RF00131: miRNA30precursor | 110 |
| RF00089: SNORD31 | 19 |
| RF00015: U4 Spliceosomal RNA | 102 |
| RF00051: miRNA17precursor | 110 |
| RF00153: SNORD62 | 53 |
| RF00020: U5 Spliceosomal RNA | 160 |
| RF00073: miRNA156precursor | 176 |
| RF00056: SNORA71 | 66 |
| RF00002: 8SrRNA | 57 |
| RF00047: miRNA2precursor | 107 |
| RF00106: RNAI | 216 |
| RF00090: SNORA74 | 210 |
| RF00066: U7snRNA | 64 |
| RF00144: miRNA199precursor | 110 |
| RF00007: U12 Spliceosomal RNA | 256 |
| RF00030: RNaseMRP | 187 |
| RF00074: miRNA29precursor | 109 |
| RF00178: miRNA24precursor | 110 |
| RF00012: U3SnoRNA | 299 |
| RF00229: IRES | 224 |
| RF00009: RNaseP | 322 |
| RF00174: Cobalaminswitch | 238 |
| RF00129: miRNA103precursor | 110 |
| RF00003: U1Spliceosomal | 170 |
| RF00008: Hammerhead Ribozyme | 98 |
| RF00004: U2Spliceosomal | 271 |
| RF00076: miRNA181precursor | 110 |
| RF00138: SNORD16 | 34 |
| RF00092: SNORA63 | 130 |
| RF00005: tRNA | 106 |
| RF00019: YRNA | 187 |
| RF00150: SNORD42 | 1 |
| RF00001: 5SrRNA | 152 |

**Table S3:** Randomly selected proteins from the *D. melanogaster* annotation used in training of the coding sequence classifier. The list below contains all those that were processed using the `Svhip` pipeline that produced feature vectors used in training.

| protein name | # training instances | # sequences |
| --- | --- | --- |
| AttC | 174 | 97 |
| Fgop2 | 129 | 116 |
| Cyp6a19 | 248 | 29 |
| Idgf5 | 264 | 116 |
| MagR | 14 | 116 |
| Alg10 | 252 | 113 |
| Bro | 73 | 115 |
| Ercc1 | 59 | 31 |
| Capa | 220 | 95 |
| Atg3 | 383 | 41 |
| Membrin | 151 | 116 |
| Crk | 91 | 57 |
| CapaR | 278 | 66 |
| mat | 77 | 114 |
| dgt2 | 66 | 116 |
| Arc1 | 117 | 114 |
| Gr85a | 207 | 93 |
| AQP | 148 | 117 |
| eRF1 | 247 | 117 |
| Fibp | 169 | 108 |
| MED10 | 53 | 117 |
| List | 109 | 20 |
| Hmgcl | 68 | 13 |
| Ing3 | 103 | 14 |
| CBP | 45 | 16 |
| Drip | 97 | 115 |
| gammaCOP | 413 | 53 |
| CSN6 | 146 | 117 |

**Figure S1:**
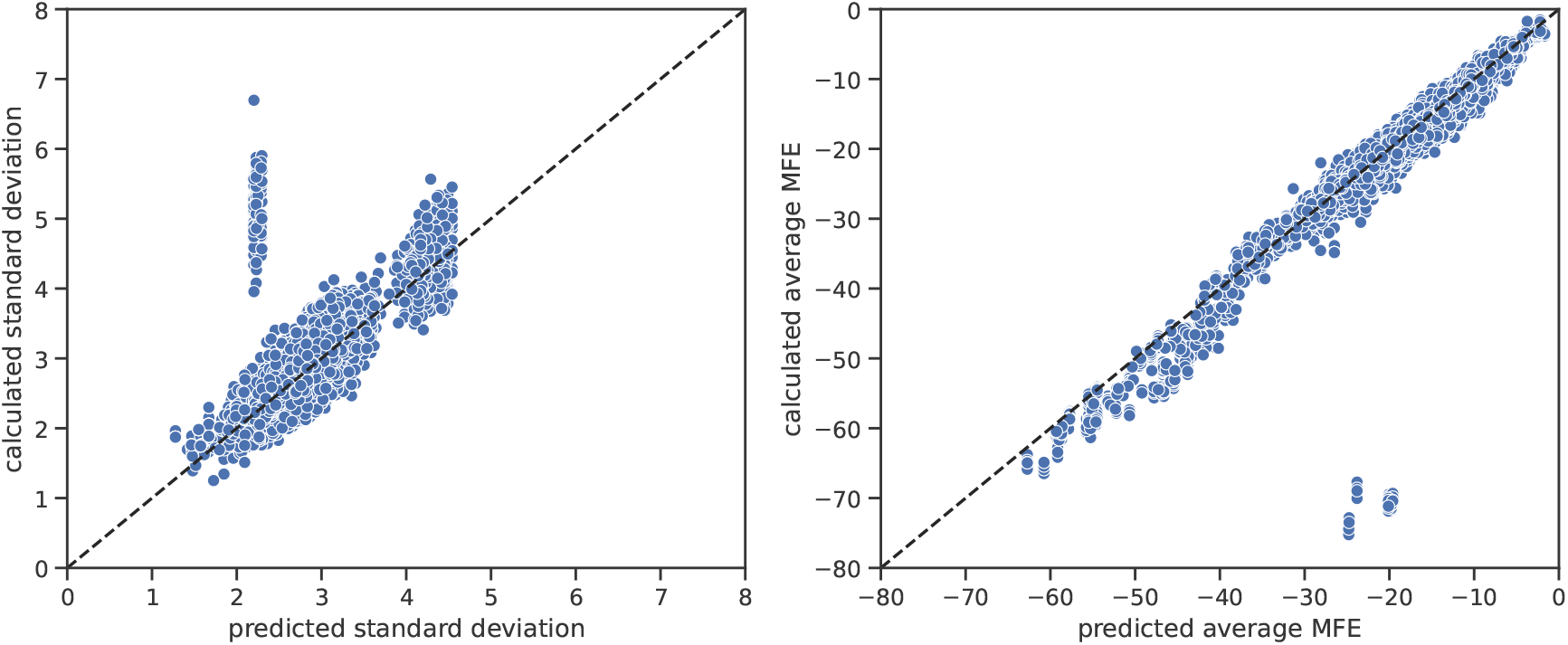
Individual predictions of average MFE and standard deviation of our SVR model. Values are used for the prediction of the alignment-wide mean *z*-score of MFE. Data points correspond to the predicted values in Figure 3.

**Figure S2:**
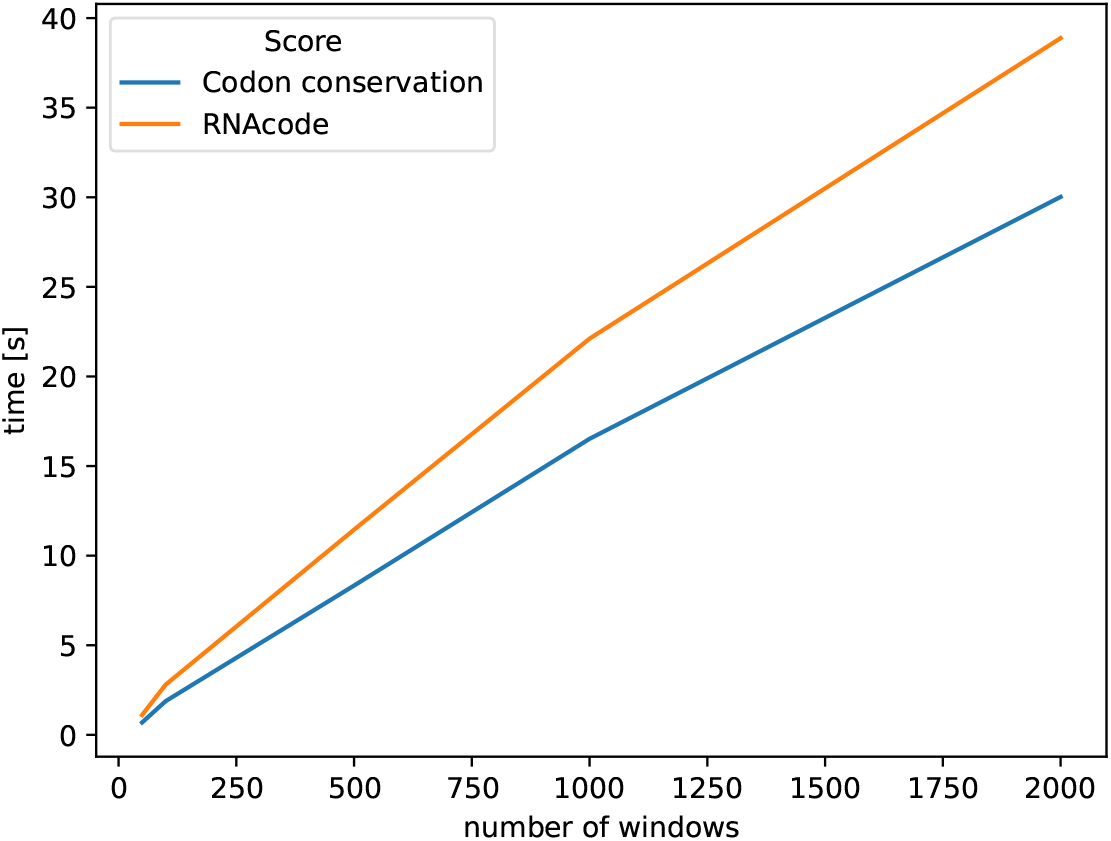
Comparison of the run time between `RNAcode` using only the best hit per alignment and the calculation of the codon conservation score. The first 2000 alignment windows were extracted from the Y chromosome MAF alignment of the *Drosophila* genome, and run time was measured on a continuous basis. The trend of the CCS outperforming the `RNAcode` score in the long run due to its lower computational complexity is clearly visible.

**Figure S3:**
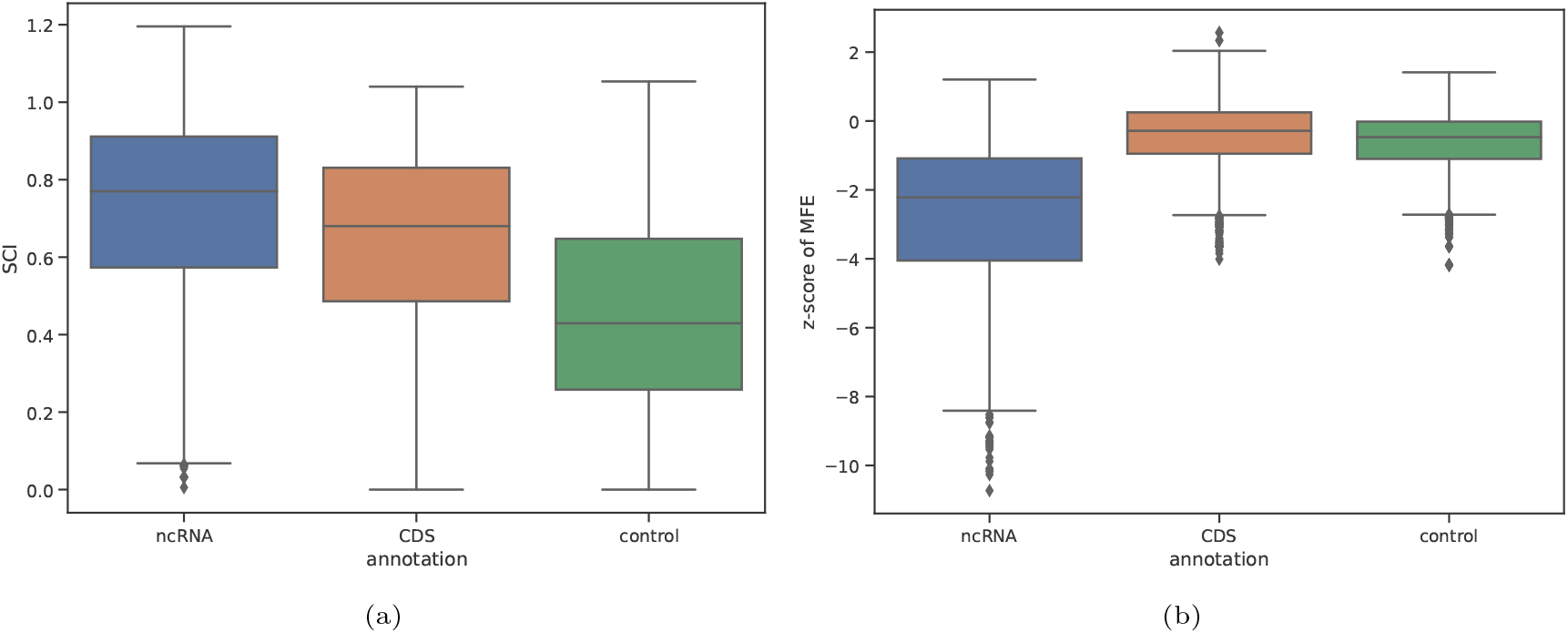
Distribution of secondary structure conservation features in protein-coding sequences when compared with non-coding RNA and an ambiguous background. The training data for the generation of the protein coding SVM model as described in Section 2.1 was used. Substantial differences between the SCI in coding sequences and the control can be observed. This does not hold for the *z*-score of MFE, which yields very similar values.

**Figure S4:**
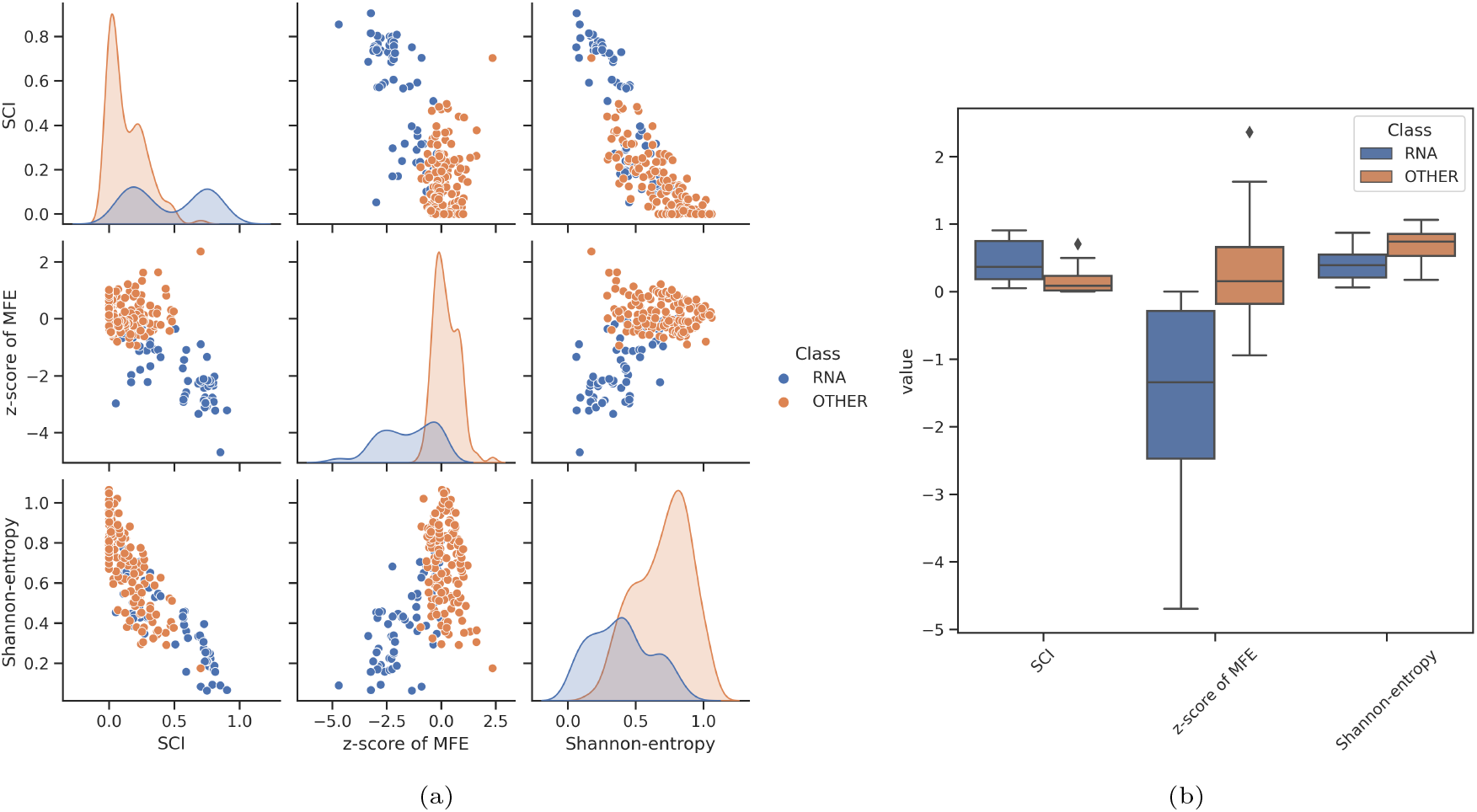
Graphical output automatically generated by `Svhip` for the alignment of bacterial RNAse P (Rfam ID: RF00010) RNA sequences. For an explanation of the output, refer to Figure 6. It can be assumed that the bimodal form of the RNA distribution and the strong partial overlap with the control group is partially caused by the presence of both prokaryotic and eukaryotic sequences in the initial input file, resulting in suboptimal alignments for a non-negligible number of alignment windows. In contrast to only the bacterial RNAse P alignment (RF00009), the class separation is less obvious in this particular case. In general, this figure serves to illustrate that using suboptimal alignment data as initial input (i. e. alignments of genetically very different species) may result in the selection of suboptimal alignment windows and thus potentially in less clear class separation.

## Footnotes

1 The annotation version used in this work is denoted as *dmelr*6.44 – *FB* – 2022

2 http://www.bioinf.uni-leipzig.de/supplements/22-003

3 https://www.uni-leipzig.de/forschung/wissenschaftliche-laufbahn/promotion/pre-doc-award

